# Omnidirectional and omnifunctional connectivity analyses with a diverse species pool

**DOI:** 10.1101/2020.02.03.932095

**Authors:** Daphnée Lecours Tessier, Roxane Maranger, Timothée Poisot

## Abstract

Connectivity among habitat patches in both natural and disturbed landscapes needs to be accounted for in conservation planning for biodiversity maintenance. Yet methods to assess connectivity are often limited, because simulating the dispersal of many species is computationally prohibitive, and current simulations make simplifying assumptions about movement that are potentially erroneous. Here we show how these limits can be circumvented and propose a novel framework for the assessment of omnifunctional and omnidirectional connectivity in a 28000 km2 area in the Laurentian region of Québec, Canada. Our approach relies on (i) the use of *Omniscape*, an improved version of *Circuitscape* which allows omnidirectional simulations that better emulate animal movement and (ii) the synthesis of large volume of species-level dispersal simulations through *a posteriori* clustering of the current intensity. Our analysis reveals that the movement of 93 species evaluated can be clustered into three functional dispersal guilds, corresponding to mostly aquatic species, terrestrial species able to use aquatic environments, and strictly terrestrial species. These functional guilds do not share connectivity hotspots, suggesting that corridor planning would need to account for the multiplicity of dispersal strategies. Although this approach requires a large volume of computing resources, it provides richer information on which landscape features are critical to maintain or need to be regenerated for broader biodiversity maintenance goals.

## Introduction

Natural ecosystems are facing unprecedented threats as a function of both direct and indirect anthropogenic disturbances, leading to a global and accelerating decline in biodiversity (Hooper et al., 2012; Merino et al., 2019). Some of the main threats to biodiversity maintenance are the joint effect of reduced area of natural habitat and connectivity loss among different habitat patches across the landscape (Thompson et al., 2017). It reduces population size, gene flow, and diversity, which in turn can lead to inbreeding and local extinction (Jackson and Fahrig, 2011). It can also prevent species from reaching new suitable habitats, and therefore limits their chance of both survival (Smith et al., 2013) and adaptation to climate change (Nuñez et al., 2013). A possible solution is to establish ecological corridors between protected areas (Rayfield et al., 2011), or more broadly to facilitate movements across the landscape at large spatial scales (McRae Brad H. et al., 2008). As ecosystems are home to multiple species with different needs and behaviours, it is essential that these corridors be multifunctional, *i.e*. that they account for the fact that the ability to move across a landscape is perceived differently by different species (Albert et al., 2017). Corridors that are well planned and implemented have the potential to promote survival, migration, and preserve small populations, for a large number of species (Smith et al., 2013). Indeed, they are essential for successful biodiversity maintenance in the long-term.

In order to establish new ecological corridors, several methods to assess connectivity have been developed in recent years. Belote et al. (2016) selected corridors with a least-cost path analysis across the continental US, using Linkage-mapper (McRae et al., 2011). Cote et al. (2009) used graph theory to assess structural connectivity of the Big Brook river drainage network (Terra Nova National Park, Newfoundland, Canada): lakes were nodes linked by river reaches. Finally, McRae et al. (2008), created Circuitscape, which used circuit theory to quantify the possibility of crossing each territorial patch and tested this for Caspers Wilderness Park in California. Despite a large variety of accessible techniques, most studies focused on the needs of terrestrial species only (Appendix 1). Indeed, according to Correa Ayram et al. (2016), only 7% of connectivity studies were conducted on fluvial habitats, while terrestrial ecosystems represent 88% of research efforts on this topic between 2000 and 2013. However, without proper consideration of connectivity to freshwater habitats (lakes and rivers), downstream health of river reaches could degrade and critical riparian wetlands could dry up, resulting in both terrestrial and freshwater biodiversity loss (Herbert et al., 2010; Smith et al., 2013). These ecosystems are naturally integrated through their watersheds (Ballinger and Lake, 2006), therefore we argue that this is the most ecologically relevant unit in which connectivity should be assessed.

To evaluate landscape connectivity for entire communities, it has been suggested to analyse a territory’s features according to the needs of a diverse number of terrestrial and aquatic species (Albert et al., 2017; Sahraoui et al., 2017). It should lead to a more robust and realistic output, as a small number of focal species seldom encompass the whole range of dispersal behaviours (Liu et al., 2018; Marrotte et al., 2017; Meurant et al., 2018). However, because multi-species analyses are complex (Moilanen et al., 2005), most efforts have often relied on the movement of a limited number of species (see Appendix 1). Moreover, according to Correa Ayram et al. (2016), there is a great redundancy of the species selected for the connectivity analyzes.

Mammals alone would represent 41% of the analyzes, mammal carnivores more than 20%. One of the issues with assessing connectivity across multiple types of habitats is that it requires to merge the results of several species-level connectivity analyses, to account for the variety of ways in which different species react to different types of landscape elements (Albert et al., 2017). When done with a large number of organisms, this imposes a double burden. One is to reconcile the connectivity of possibly dozens of indicator species, not all of which may have unbiased habitat use models (Olden et al., 2002). The other is having the computational capacity to actually carry out this analysis. As the later is becoming less of a problem with increased computational power and more diversified software offering to carry out connectivity simulations, developing a general framework to include connectivity across ecosystem types, with a diverse set of species, is an achievable and timely task.

A few studies have recently developed methods to aggregate multiple species-specific analyses in a single result. For example, Albert et al. (2017) and Sahraoui et al. (2017) selected 14 and 16 eco-profiles respectively, each composed of one or more species, in order to represent a broader range of dispersal behaviours and habitat requirements. These eco-profiles were designed through dimensionality reduction based on species connectivity and habitat needs, either using clustering or multivariate analysis. In another effort, Santini et al. (2016) selected all the 20 non-volant mammal species of Italy that had a dispersal ability of a median of 3 km to identifying defragmentation priorities. The three studies have created composite maps by summing the single-species results or by calculating the mean score for each patch of habitat. Santini et al (2016) went even further by testing seven other ways to merge the species-specific maps, using weighted means of different landscape features to assess link probabilities, and two different summation types for nodes in the network. These studies have demonstrated that it is possible to create connectivity analyses for a large number of species, which may be ecologically more relevant. Indeed the results could be meaningful for decision-makers, helping them to target new protection areas, restore others, and inform more sustainable strategies in city development projects (Ersoy et al., 2019). Despite these successes, they remain limited by three key aspects. First, even a pool of 20 species cannot represent the entire functional diversity of realistic communities. Second, selected species pools continue to under value the contribution of semi-aquatic species and their habitats to landscape connectivity. Furthermore, freshwater biodiversity is considered the most imperiled globally (Hendriks, 2016), suggesting that the need to include aquatic and semi-aquatic species is urgent. Finally, because species clustering is done before the simulations, this assumes that species connectivity depends on habitat requirements alone and neglects differences that could emerge due to landscape configuration.

In this study we present a general framework for multi-species connectivity analysis, built around the needs to (i) integrate terrestrial and aquatic ecosystems (Herbert et al., 2010), (ii) use an ecologically relevant number of species to capture the diversity of ways in which species interact with their landscape (Meurant et al., 2018), and (iii) provide measures of uncertainty after different analyses have been merged. By contrast to previous studies relying on *a priori* clustering of dispersal guilds (Lechner et al., 2017; Sahraoui et al., 2017) we propose an *a posteriori* method for the creation of composite indicator species-guild. Our approach accounts for the way species are predicted to move across the landscape by looking for similarities in their dispersal simulations, rather than in the cost matrix of these species. We argue that this new way of interpreting and post-processing connectivity simulations will result in a more integrative view of landscape connectivity to favour biodiversity maintenance in the long-term.

## Methods

### Study area

We assessed our new connectivity approach using a 27 994 km^2^ area located on the north shore of the St. Lawrence River, near the city of Montréal, Québec, Canada (Fig. 1). Seven major rivers are included in this region, and the area sits on two strikingly different geological provinces, the Saint Lawrence lowlands, dominated by agriculture and urban areas, and the Canadian Shield, covered largely by pristine temperate forests with a very high density of lakes and rivers (19.8%). This high aquatic surface supports the need to account for more semi-aquatic species in our simulations. This region was chosen because it shows a strong gradient of anthropogenic disturbance, with heavy urbanization and agricultural development to the south, major highways on the north-south axis, many roads, large forested areas particularly on the Shield where there is an active forestry industry, some major National Parks (Mont Tremblant, Oka, la Mauricie), and several small cities that service a large cottage industry. Moreover, biological surveys conducted by conservation agencies and different government bodies resulted in the creation of a curated list of species, which we integrated in this analysis.

**Figure 1.**
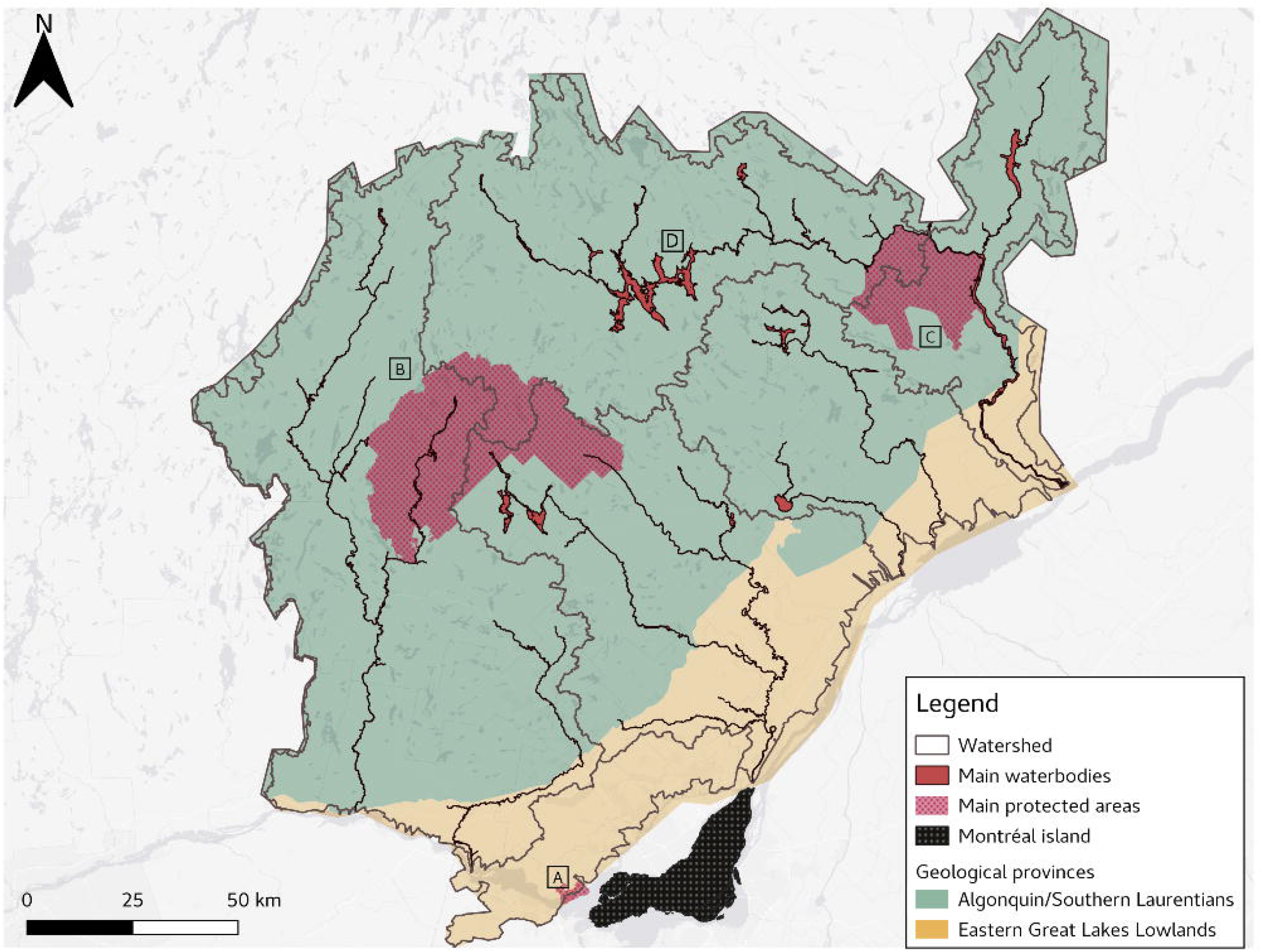
Study area, on the North Shore of the Saint Lawrence river, province of Québec, Canada. A. Oka National Park B. Mont-Tremblant National Park C. La Mauricie National Park D. Lac Taureau. The study area is astride two geological provinces, the Saint Lawrence Lowlands (yellow) and the Canadian Shield (green).

**Figure 2.**
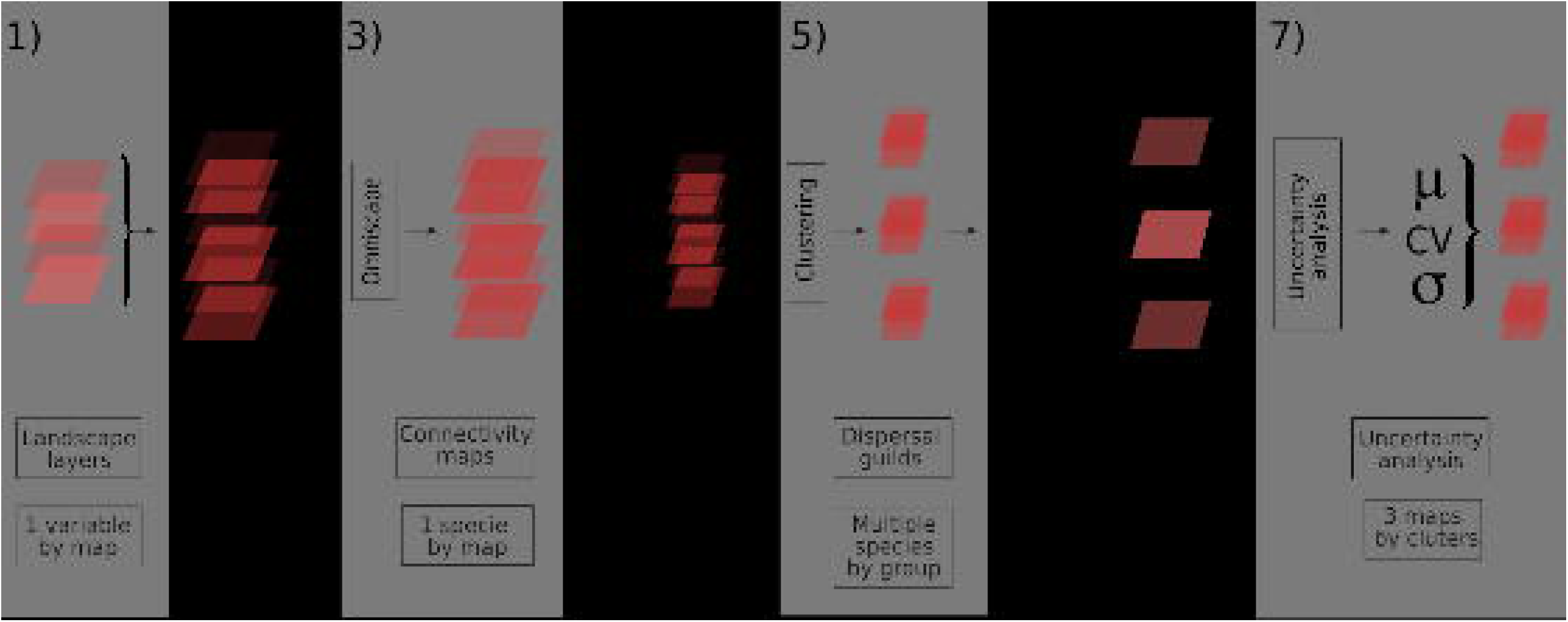
Representation of the overall method flow. For each species (93), landscape layers features got scored between 0 and 100 for their resistance (1). Then, the layers were converted in a raster format to be summed, which created one resistance map by species (2). Following this step, the maps were analysed using Omniscape, which created connectivity maps (3). We assigned to each pixel its quantile in the observed map of the cumulative current for the species (4) and conducted a clustering fuzzy C-means analysis on the unfolded 93 connectivity map to get the *a posteriori* dispersal guilds (5). After the clustering, we extracted the weighted centroids of each cluster (6) and measured the additional indices (evenness, variation, location of the most connected pixels) (7).

#### Selection of species

In order to select the species for this study, two lists of recorded species from the main National Parks were used (SEPAQa, 2019; SEPAQb, 2019). We kept 45 mammals, all amphibians (16) and 9 reptile species, plus 15 avian species with a conservation status. We added five more avian species, two amphibians and one reptile, which have been recorded in GBIF and Ebird database since 2000, for a final list containing 93 species (see Appendix 2).

#### Collecting species habitat preference data

From the list of focal species, we got information on habitat preference and the way they move across the landscape. Since these data are very heterogeneous, we gathered them from several sources. We used data from the IUCN Red list (IUCN, 2019.), identification guides of Quebec fauna (Desroches and Rodrigue, 2018; Prescott and Richard, 2014) and expert-curated websites for the herpetofauna (“aarq | EcoMuseum,” 201.) and the avian fauna (Écopains d’abord, 2019.) The complete cost matrix for all species is given in **APPENDIX 2**.

#### Creating the resistance maps

**Resistance maps** are a mix between energetic landscape and habitat quality modelling. They represent the energetic and ecological cost to cross a certain plot as a function of its configuration (topography, human structures) and composition (land cover) (Table I) (McRae Brad H. et al., 2008). Matching preference data with the information on the land cover and human structures present in the area allowed us to create a landscape resistance canvas for every species, where 0 represents the absence of resistance and 100 the most important obstacle to dispersal. Resistance maps are the sums of every spatial feature that could influence a species dispersal.

According to the resistance canvas, we created the different layers composing the final resistance map, using Qgis 3.4 (Qgis Development Team, 2019). We began with the land cover layer, formed by thousands of polygons of 18 habitat types (agglomeration, crops, coniferous forest, herbaceous wetland, etc.). To transform it into a resistance layer, we integrated the resistance score of each species, to each type of land cover in the layer’s data (Appendix 2). This is the basic landscape features which influence species dispersal. In order to assess landscape complexity to its fullest extent, we added the effect of slope, buildings presence, waterbodies, roads, trails, and railways (Appendix 2) on species dispersal. For the roads, we have accounted for their effect on a broader scale (150 m), with the resistance score decreasing according to the distance from the road (Forman and Alexander, 1998). For the “slope” layer, we have considered the degree of the slope as the degree of resistance (a slope of 50 degree equals a resistance of 50). Indeed, slope is known to have an impact on species displacement movement (Gaudry et al., 2015; Leblond et al., 2010).

In order to create a resistance map by summing the layers for each species, we converted vector resistance layers to raster layers. We first chose to use square cells at a resolution of 2 meters^2^, which represents the smallest disturbance width considered in this study, hiking trails. Then, for each individual species, we summed the resistance layers corresponding to their canvas, which produced the 93 species-specific resistance maps. The resistance score of each cell represents the sum of all landscape features, according to the species dispersal behaviour canvas.

**Table 1.**
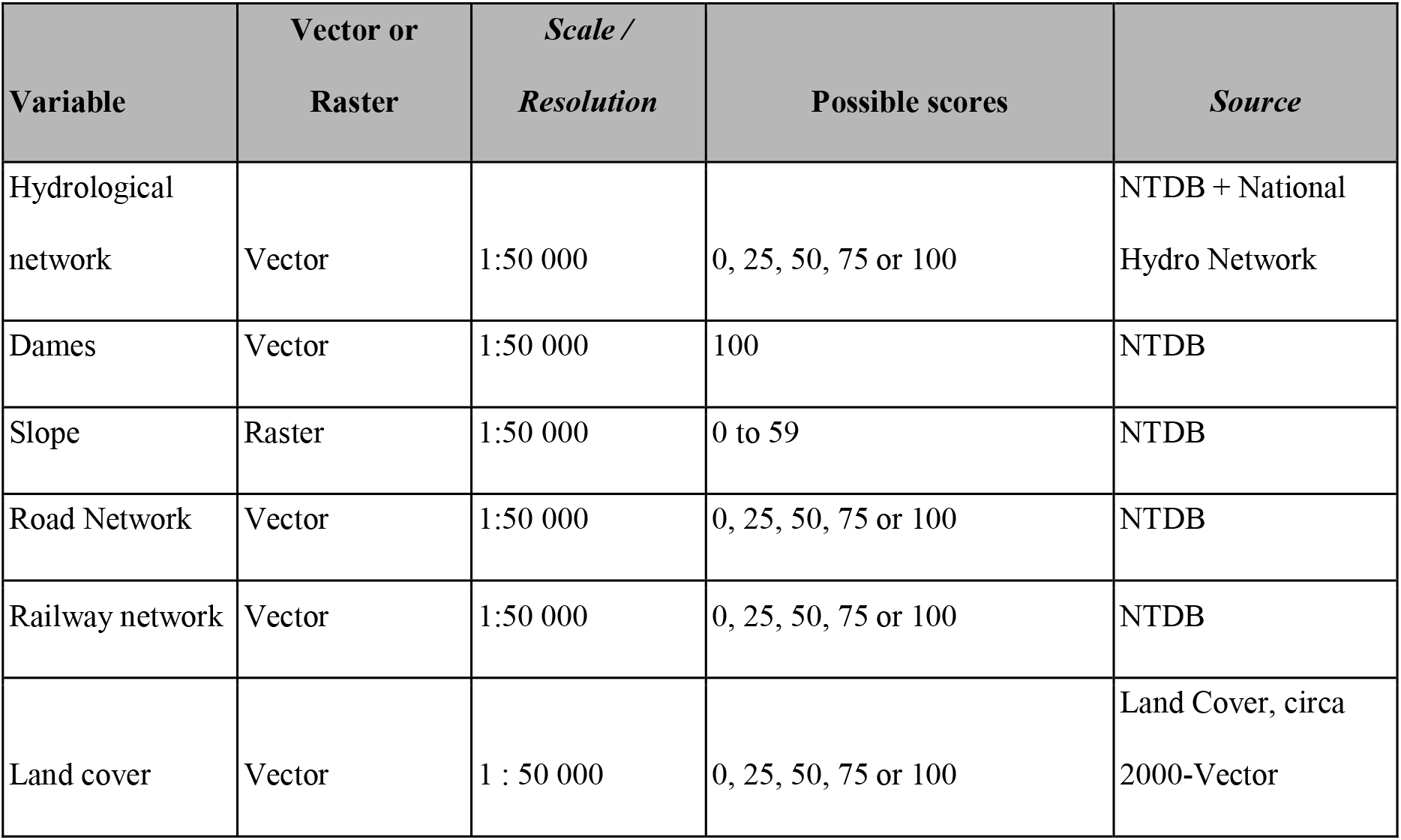
Compilation of the data use, their type, resolution, the possible scores attributed to and their source

#### Analyzing landscape connectivity using Omniscape

We analysed species-specific resistance maps in *OmniScape* 0.1.2 (McRae et al., 2016), implemented with julia 1.2.0 (Bezanson et al., 2015). *OmniScape* uses a round sliding windows approach to connectivity simulations, which amounts to assuming that the species is aware of its immediate surroundings, but not of the entire landscape. Indeed, species use the landscape without knowing it entirely, which is assumed by the least-cost analysis (McRae Brad H. et al., 2008). We used three different radiuses (30, 60 or 120 blocks, each block being a square parcel of 180m^2^), and averaged the quantiles of the cumulative current map for all. This resulted in connectivity scores ranging from close to 0 (specifically the inverse of the number of simulated pixels) to 1. The choice to examine quantiles as opposed to raw cumulative current values was driven by the fact that although different species will disperse at different rates (and therefore will have different total cumulative currents), quantiles provide information as to their relative use of the landscape. This would therefore facilitate the search for functional dispersal guilds. Note that for some applications, analyzing the normalized current (*i.e*. the cumulative current divided by the potential flow, as in *e.g*. McRae (2008), which is to say measuring how *surprising* the observed current is compared to a null expectation) is preferable, notably for corridor mapping or the identification of dispersal bottlenecks. In this analysis, we focus on how organisms could disperse in the landscape, and therefore use the cumulative maps.

All simulations were performed on the beluga supercomputer, operated by Calcul Québec; the project required an estimated total of 3 core-years to complete, being roughly equivalent to 3 years of time on a single-core machine.

#### A posteriori functional dispersal guilds

After all the simulations were complete, we aggregated the maps using fuzzy C-means (C. Bezdek, 1981; Dunn, 1973). This approach allows to have species contribute to more than a single cluster (for example, species that are terrestrial but have an affinity for water will have landscape use that borrows from strictly aquatic and strictly terrestrial species), as captured by the fuzziness parameter. Following community-established best practices, we set this parameter to 2. Specifically, we applied fuzzy C-means by transforming all the connectivity scores into a *s* by *p* matrix, with *s* the number of species and *p* the number of pixels with non-null connectivity values, then by calculating the *s* by *s* Euclidean distance matrix between all pairs of species. We extracted the centers of every cluster, and the species with the highest weight was the most representative species for the dispersal guild. It may therefore be used as an indicator species in the future. In addition, the weight of each species for each cluster can be used to measure species specificity; we used PDI (Poisot et al., 2012) as a specificity estimator, which has the desirable property of returning values that are lower than 0.5 for generalist species, and values above 0.5 for specialist species.

Fuzzy C-means lacks an established way to optimize the number of clusters[12]. As our goal is to identify functional dispersal guilds that move across the landscape in different ways, we relied on a simple heuristic to identify the optimal grouping. Fuzzy C-means results in a *c* by *s* matrix, where *c* is the number of clusters; every column in this matrix is therefore the contribution of all species to a cluster. Based on this information, we computed a *c* by *c* correlation matrix between clusters, and retained the solution in which no pair of clusters had a positive significant correlation.

#### Uncertainty analysis

To produce the final connectivity map, we averaged the scores for the centroids of the clusters identified in the previous step, measured the standard deviation within each pixel, and finally measured the coefficient of variation and Pielou’s evenness within each pixel. The average connectivity gives an idea of dispersal potential, and the coefficient of variation and evenness informs about the areas in which the guilds differ markedly in their landscape use. In addition, in keeping with the Aichi objectives (SCBD, 2011), we mapped the best 17% of pixels (*i.e*. with the highest connectivity scores), using the average connectivity map as well as the map for every dispersal guild.

## Results

### Guild clustering

From the 93 connectivity maps created, the fuzzy C-means identified an optimal fuzzy partitioning into three functional dispersal guilds (FDG henceforth), whose most representative species are the <u>Horned grebe</u> (*Podiceps auratus*, Linnaeus, 1758) (FDG1; fig. 3a), (FDG2; fig. 3b), the <u>American black bear</u> (*Ursus americanus, Linnaeus, 1758)* and <u>Northern Two-lined Salamander</u> (*Eurycea bislineata*, Green, 1818) (FDG3; fig. 3c). All of these guilds share a very poor connectivity score in the lowlands (although FDG3 can use some of the wooded areas for east-west traversal), and differ in how they use the rest of the study area. In particular, FDG3 tends to have almost unrestricted movement outside of the lowlands, while FDG2 is predicted to be more canalized in some places. Interestingly, while the two guilds for the more terrestrial species were not correlated (rho = 0.054), they were both correlated negatively to the cluster with more aquatic species (rho approx. −0.7 in both cases). This reveals not only that the needs of aquatic and semi-aquatic species are different, but that favoring terrestrial species only in connectivity planning will impede the movement of semi-aquatic species. The complete clustering result, including specificity analysis, is given in **Appendix 3**. Applying PDI on the clustering weight identified 15 generalist species, including notably the Canada Lynx (*Lynx canadensis*, Kerr, 1792) (PDI = 0.45), moose (*Alces Alces*, Linnaeus, 1758), (PDI = 0.44), Red-headed woodpecker (*Melanerpes erythrocephal*us, Linnaeus, 1758) (PDI = 0.27), and Hoary bat (*Aeorestes cinereus*, Beauvois, 1796) (PDI = 0.42). Whereas specialist species, or species with high affinity for a specific FDG can be good indicator species, generalist species can provide an overall view of landscape connectivity.

**Figure 3.**
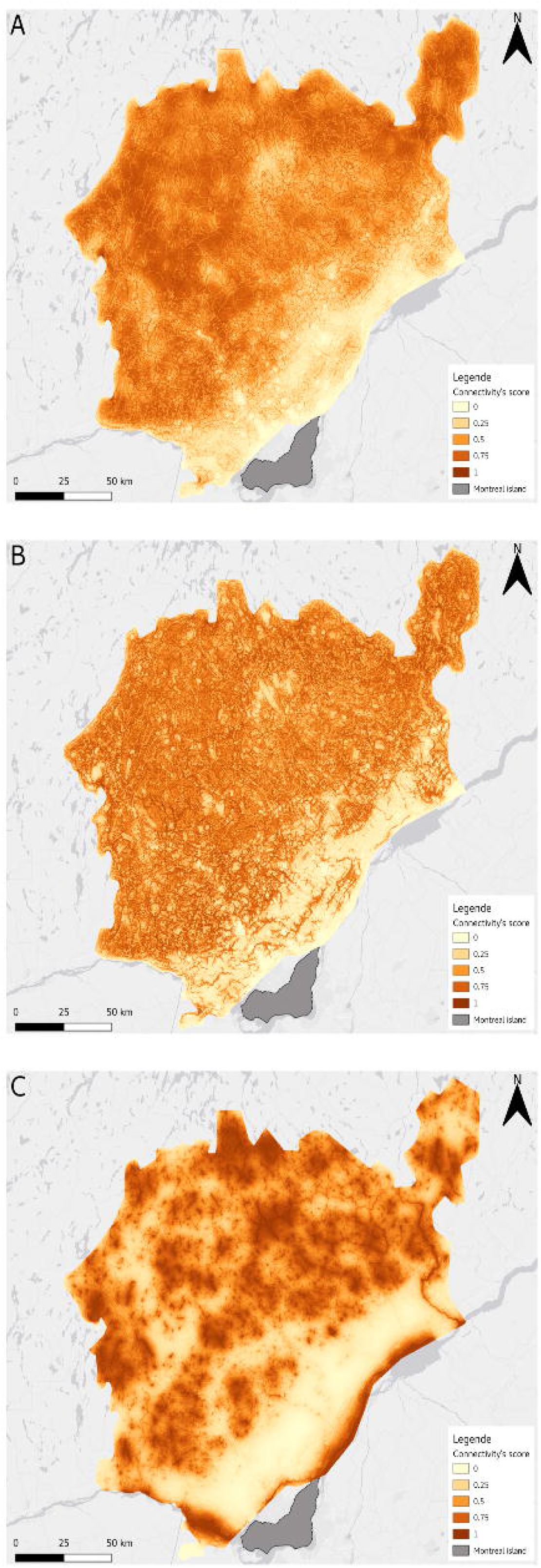
Connectivity scores for the three different guilds. Higher values indicate more potential for movement. St. Lawrence Lowlands have areas with medium connectivity potential for FDG1 and FDG2 but are globally lowly connected and do not have real potential to facilitate the movement of species from FDG3. The Canadian shield, however, show high connectivity potential for the three guilds, especially FDG2. A. FDG1 – Species of open and undisturbed habitats B. FDG2 – Species of forested habitats C. FDG3 – Species of freshwater habitats

### Average connectivity and variation

As expected, the St-Lawrence lowlands performed uniformly worse in terms of connectivity, with nevertheless some areas running parallel to the shore that can work as dispersal avenues (fig 3a), though these are mostly used by FDG3. Even in the more forested areas it is strikingly easy to identify more developed regions or axes, corresponding to highways connecting the island of Montréal to smaller clusters of anthropogenic activity. With the exception of these areas, the region is remarkably well connected overall. The coefficient of variation and Pielou’s evenness for connectivity reveals areas in which the needs of species that favour either aquatic or terrestrial habitats are more difficult to reconcile (fig 4b, 4c). This includes, trivially, the St Lawrence river itself, but also areas surrounding large water bodies such as the *Lac Taureau* to the North. It should be noted that although both the coefficient of variation and Pielou’s evenness provide a similar information (namely, the homogeneity in the use of every pixel by the three FDGs), the later is a more appropriate measure. Although widely used, the coefficient of variation performs better on ratio scales, whereas the quantiles of dispersal, being bounded, are expressed on an interval scale.

**Figure 4.**
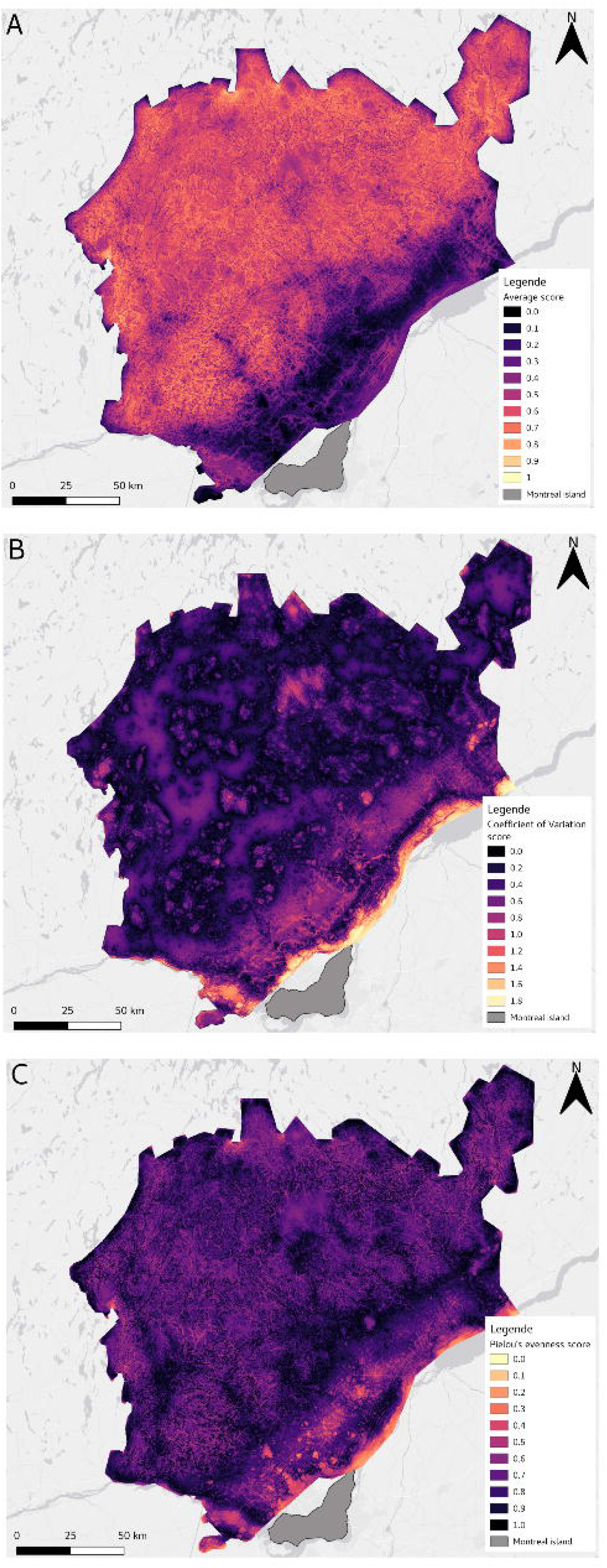
Representation of the Average connectivity score (A), Coefficient of variation (B) and Pielou’s evenness (C) A. Average connectivity across the three functional guilds. The St. Lawrence Lowlands have a lower connectivity overall, though some more intact habitats running parallel to the shore allow for traversal. B. Coefficient of Variation across the three FDGs (n=3), corrected for small sample size by a coefficient of (1+1/4n). Values on the map have been capped at 1, as values larger than unity are generally regarded as noise. C. Pielou’s evenness across FDGs. Note that the color scale has been inverted, to reflect the fact that high values correspond to more similar connectivity across FDGS.

### Best-connected pixels

In fig. 5a, we represent the 17% of pixels with the highest *average* connectivity, which can be thought of as forming the basis for a dispersal network across the region, independent of guilds. This information should be contrasted to fig. 4b, in which we show the spatial distribution of the best 17% of pixels for each of the different FDGs and superimposed them. Because the guilds are optimized to maximize the differences between them, their most connected areas rarely overlap. As a result, protecting the top 17% of pixels for all guilds combined would cover up to three times more surface than the best 17% pixels on average. Furthermore, note that very few aquatic habitats would be protected using the average classification as a target.

**Figure 5.**
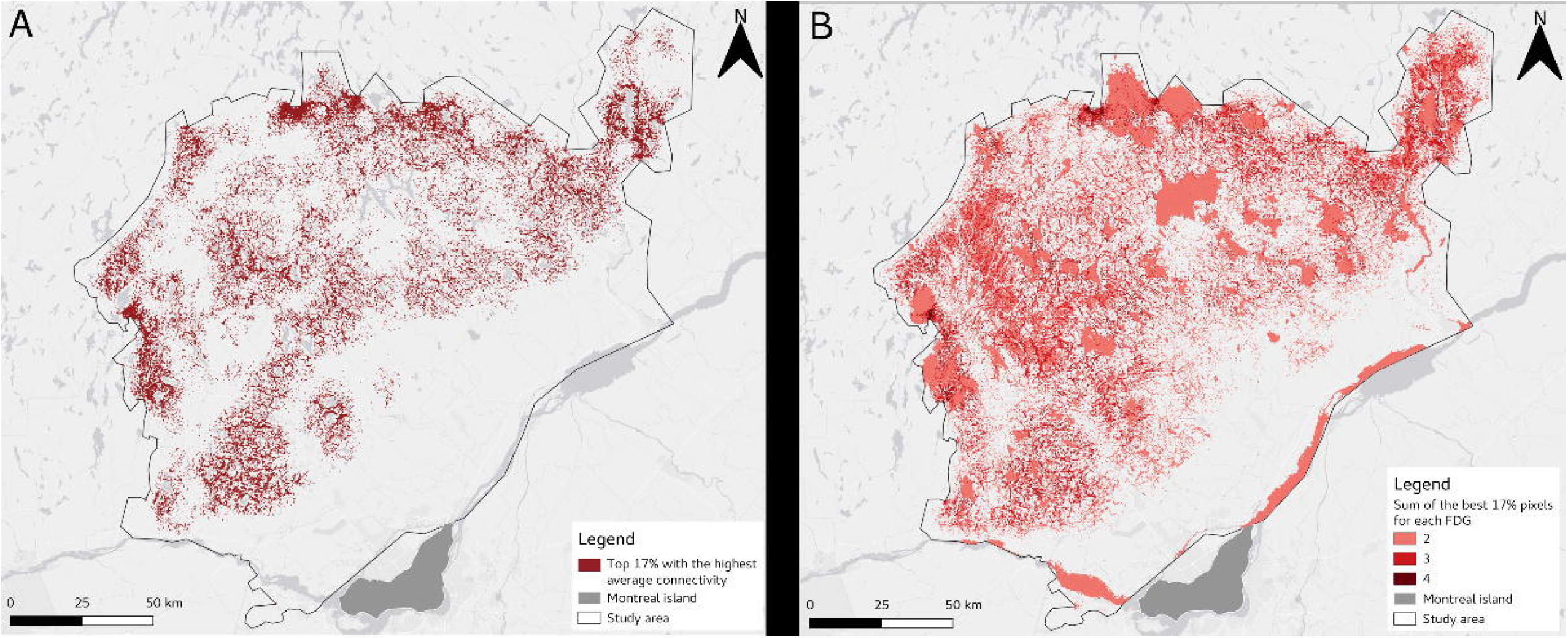
Spatial distribution of the best 17% of pixels with the highest average connectivity (A) and the best 17% of pixel for all guilds combined (B). The pixels in panel B cover approximately 40% of the entire territory, suggesting that the three functional guilds have very high complementarity.

## Discussion

We have simulated the dispersal of 93 species across a 220 by 230 km area in the Laurentian region of Québec. We then classified the multi-model average of connectivity maps using Fuzzy C-means, to provide a regional average (Fig. 4a). Based on this result, we identified areas of higher uncertainty (Fig. 4b, 4c) as well as candidate indicator species for each of the three Functional Dispersal Guilds (FDGs). We found that connectivity is lower in the St Lawrence Lowlands, which is more impacted by anthropogenic activities, and generally better in the forested areas of the Canadian Shield. Although this result was expected, our approach was the first to assess connectivity across such a large heterogeneous landscape, by accounting for the full taxonomic diversity of both terrestrial and semi-aquatic species. Furthermore, these species belong to different guilds representing three emergent dispersal landscapes that coexist within the study region. Conservation actions that may neglect the protection of a specific guild due to poor representativity of indicator species could have overall negative consequences on the biodiversity maintenance of the entire region. Indeed, semi-aquatic, which form a guild on their own, are often neglected.

We have used fuzzy C-means to cluster the species into FDGs, as it represents a more objective approach in which species are able to contribute relatively among dispersal guilds. Indeed, we identified about 16% of species classified as generalists that contribute in similar proportions to all FDGs. Under a non-fuzzy scheme, this information would be lost as species would be forced into one guild only. This being said, as we show in fig 6, the results of the fuzzy C-means algorithm are directly comparable with a hierarchical clustering using Ward’s agglomeration on the pre-squared distance matrix. Although simple hierarchical clustering would have led to the same dispersal guilds, the rich information of the movement of generalists species among guilds could only be achieved using the fuzzy C-means approach.

**Figure 6.**
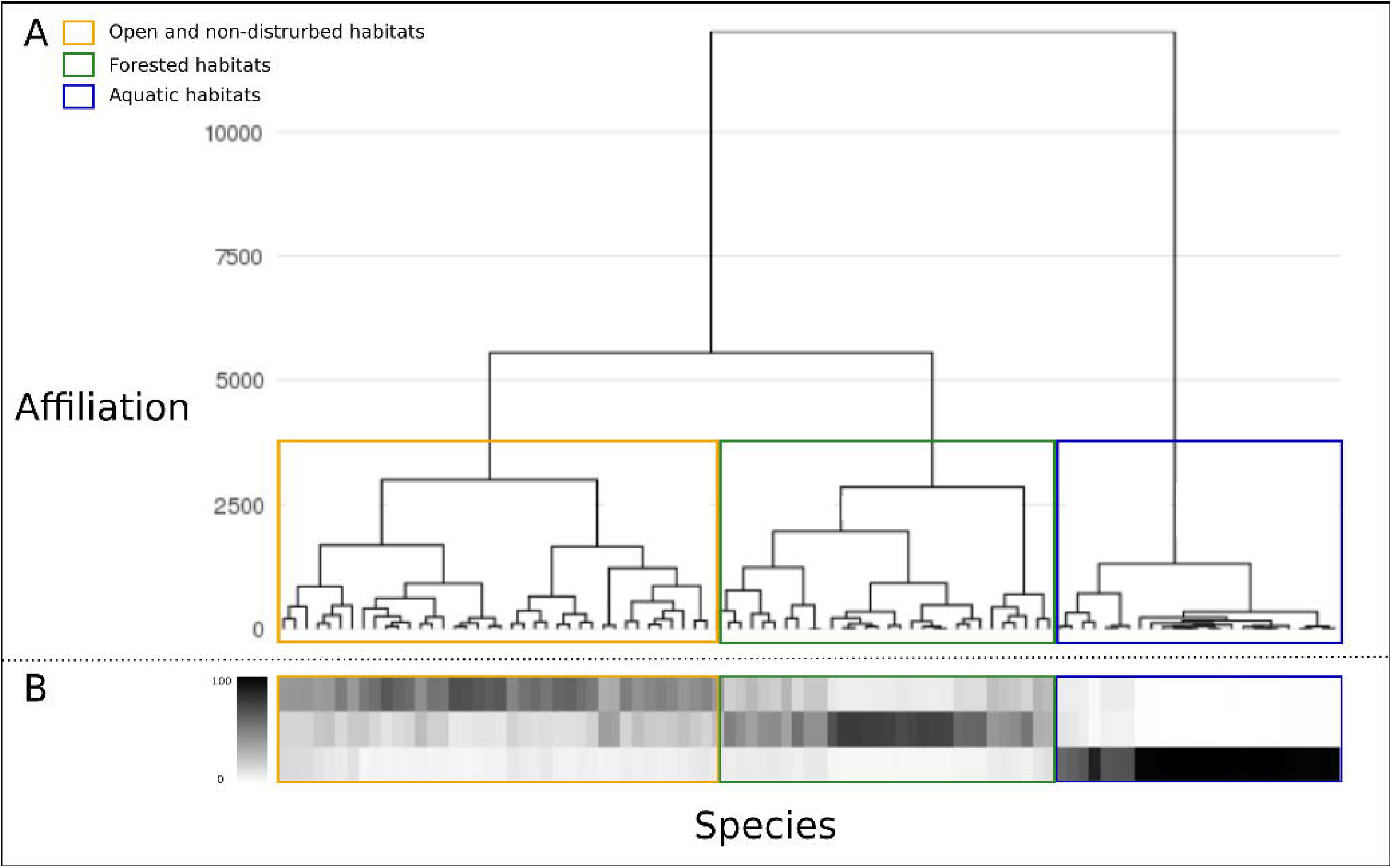
Comparison between Ward’s clustering (A) and the Fuzzy C-Means (B) results. Both methods identify three distinctives groups of dispersal behaviour (guilds), but only Fuzzy C-means allows to uncover the fact that some species are actually generalists.

Clustering returns groups of species based on predicted dispersal, and therefore represents a process to select indicator species within each guild. By contrast, the *a priori* selection of indicator species risks either underrepresenting or over representing different dispersal modes. This is a particularly concerning issue, since as we illustrate in fig 4, the identification of the most connected pixels varies greatly depending on whether the FDGs, or the average connectivity, are used. Knowing the importance of the choice of representative species, the more ecologically relevant *a posteriori* clustering should be favored, especially given that this amount of simulations is computationally feasible. Indeed, among the 93 species used in this study, several that would be considered “terrestrial” only, were found to benefit from freshwater habitats for travel and therefore clustered differently. For example, this was notably the case for the clustering of moose *(Alces alces*, Linnaeus 1758) with the common garter snake *(Thamnophis sirtalis*, Linnaeus 1758) and many avian species, among others. This counterintuitive association demonstrates that with regards to landscape connectivity, our preconceived ideas of species groupings do not apply.

Despite the large diversity of species and their habitat preferences, fuzzy C-means identified an optimal grouping of three functional dispersal guilds only. Surprisingly this is a lower number of classes than most multi-species simulations have been using, and so it calls into question whether using more species would have led to pseudo-replication, or to over-representing the needs of some groups of organisms. Previous methods to suggest surrogate species for connectivity modeling also resulted in a larger number of species (*e.g*. between 5 to 7 for Meurant et al. 2018). We argue that the number of species guilds should not be determined *a priori*, since these classifications would rely only on between-species differences in habitat preferences, and would not be able to account for how these differences interact with the spatial configuration and habitat diversity in the landscape. Therefore we suggest our approach be used with the largest number of species available, particularly in biophysically complex and species rich landscapes.

Once initial clustering is made, the entire species list may no longer be required for subsequent simulations carried out for the same region. In a context such as a predicting the effect of land-use or climate change on connectivity, which would involve many thousands of simulations, one strategy would be to identify the most representative species for the optimal number of FDGs, and then use these indicator species in the simulations. While this results in the loss of some information, this offers a reasonable compromise between the effort to identify relevant species in initial computationally intensive analysis, and the effort to perform projections of future connectivity maps. Additional schemes to pick a subset of species to work on should be evaluated on a case by case basis and can involve picking a mix of specialist and generalist species.

Here we provide, to our knowledge, the most robust and fully integrated terrestrial and aquatic multi-dispersal connectivity assessment at a regional scale. We used a broad suite of diverse taxa in a heterogeneous landscape structured by a number of different anthropogenic disturbances across two major geological provinces. The integration of the multiple dispersal pathways provides a robust connectivity conservation plan and our approach offers a road map for future connectivity assessments. Given the ongoing biodiversity crisis, identifying and implementing corridors to maintain habitat connectivity for conservation for as many species as possible is increasingly urgent. Yet, current conservation strategies often neglect the diversity of ways in which species actually use the landscape, since they focus largely on terrestrial species. For example, as a worst case scenario, habitat patches crucial to the dispersal of a specific functional group would not be adequately protected; this connectivity loss for an entire subgroup of species could have a cascading effect with the potential to destabilize the entire ecosystem.

## Data availability

Data will be deposited on DataDryad upon acceptance.

## Acknowledgements

We acknowledge that the land described in this study is located within the traditional unceded territory of the Saint Lawrence Iroquoian, Anishinabewaki, Mohawk, Huron-Wendat, and Omàmiwininiwak nations. We have received financial support from the NSERC Discovery grants program (TP, RM), the Groupe de Recherche Interuniversitaire en Limnologie (DLT), the BIOS^2^ NSERC CREATE grant (DLT), the Canadian Foundation for Innovation John Evans fund (TP). Computations were made on the supercomputer beluga, managed by Calcul Québec and Compute Canada. The operation of this supercomputer is funded by the Canada Foundation for Innovation (CFI), the ministère de l’Économie, de la science et de l’innovation du Québec (MESI) and the Fonds de recherche du Québec – Nature et technologies (FRQ-NT). We thank Guillaume Spain for help with GIS. We thank Kimberly Hall, Vincent Landau, and Ranjan Anantharaman for assistance with OmniScape. Author contributions: DLT acquired the data, DLT and TP performed the analyses, all authors contributed to study design and writing.

